# Microfluidic enrichment of proteolytic microbial consortia from sewage sludge

**DOI:** 10.64898/2026.02.10.705040

**Authors:** Luca Potenza, Valentina Smacchia, Łukasz Drewniak, Tomasz S. Kaminski

## Abstract

Proteolytic microbial consortia are key drivers of protein hydrolysis in complex organic substrates. In anaerobic digestion systems, such as biogas production from sewage sludge, this process constitutes the initial and rate-limiting step. Despite their importance, proteolytic microorganisms remain poorly characterized due to the complexity of environmental microbiomes and the limitations of conventional cultivation and screening methods.

Here, we present a label-free microfluidic protocol for the high-throughput cultivation and characterization of proteolytic microorganisms. Single microbial cells are encapsulated in gelatine droplets and grown clonally, where proteolytic activity is detected through image-based analysis of droplet shape changes. Enrichment of individual proteolytic cultures is achieved using a separate passive microfluidic device that enables droplet sorting. Taxonomic characterization of sorted droplets by 16S rRNA gene sequencing revealed a fivefold higher number of ASVs and a more diverse array of proteolytic strains were recovered compared with conventional skim milk agar screening (SMA).

Taken together, this microfluidic workflow allows accurate and fast enrichment of proteolytic strains. Our approach contributes to a deeper understanding of proteolytic communities in sewage sludge and opens new opportunities for targeted microbial recovery in waste-to-energy applications.

**Importance:** Proteolytic microorganisms drive the initial and rate-limiting step of protein degradation in anaerobic digestion systems such as sewage sludge biogas production, yet their diversity and function remain poorly characterized due to the limitations of conventional cultivation methods. We present a label-free droplet microfluidic workflow that enables high-throughput, single-cell cultivation, functional screening, and selective enrichment of proteolytic microbes directly from complex communities. This approach substantially improves the recovery and diversity of proteolytic strains compared with traditional assays, providing a powerful tool to study hydrolytic consortia and to enhance microbial discovery for waste-to-energy and other biotechnological applications.

## Introduction

Biogas production represents a substantial component of current sustainable energy strategies, converting organic waste into methane-rich fuel and mitigating environmental pollution[1]. Sewage sludge, a ubiquitous by-product of municipal and industrial wastewater treatment, is particularly attractive as an anaerobic digestion substrate due to its abundance and high organic content[2]. However, the efficiency and stability of anaerobic digestion are frequently constrained by the hydrolysis step, the initial and rate-limiting phase during which complex polymers such as proteins, polysaccharides, and lipids are broken down into soluble monomers[3,4].

Recent research in biogas production has either highlighted the synergistic role of diverse hydrolytic microbial consortia in sludge hydrolysis[5,6] or focused primarily on the degradation of lignocellulosic biomass[3,7,8]. Nonetheless, protein degradation remains relatively underexplored, and the enrichment of proteolytic strains remains challenging; only a limited number of recent studies have begun to examine their diversity, ecological roles, and potential contributions to improving biogas production[9]. In this context, proteolytic consortia represent a critical yet poorly characterized component of sludge hydrolysis. Although hydrolysis is widely recognised as the rate-limiting step in anaerobic digestion, protein hydrolysis constitutes a distinct bottleneck in sewage sludge, as proteins are degraded later and less completely than carbohydrates, thereby constraining methane yield[10]. Accordingly, targeted enhancement of proteolytic activity has been shown to increase methane production[11].

Culture-independent approaches like 16S rRNA sequencing and metagenomics have broadened our understanding of sludge microbiome diversity. Yet, they provide only indirect evidence of enzymatic activity and fail to bridge the genotype-phenotype gap, particularly at the single-cell level, hindering our ability to link specific taxa to proteolytic functions in complex microbial samples[12]. Conventional methods for studying proteolytic communities, such as plating on selective media or cultivation in liquid media followed by spectrophotometric assays using chromogenic or fluorogenic substrates, are inherently biased toward fast-growing or easily cultivable microbes. Moreover, artificial substrates may not accurately reflect in situ activity, limiting ecological relevance and functional discovery[13]. The skim milk agar (SMA) assay is a common method for screening microbial protease activity. Protease-producing microorganisms hydrolyse casein, forming clear halos around colonies, while non-proteolytic strains leave the medium opaque. Halo size relative to colony growth provides a semi-quantitative proteolytic index. SMA is inexpensive, simple, and allows rapid visual discrimination of protease-positive strains in both pure cultures and mixed communities. However, halo formation depends on growth rate, colony morphology, and medium diffusion, potentially biasing results, and it only offers qualitative or semi-quantitative information without capturing full protease specificity or environmental activity. In addition, non-proteolytic mechanisms such as acidification can occasionally alter medium opacity, leading to false positives[14]. Another popular method is represented by gelatine-based assays that detect protease activity as protease-secreting microbes hydrolyse gelatine, breaking down the gel and causing medium liquefaction. Gelatine serves as both a nutrient and a gelling substrate, and liquefaction provides a simple qualitative measure of proteolytic activity.

Droplet microfluidics offers a powerful alternative to traditional cultivation by encapsulating individual microbial cells in millions of picoliter-scale droplets that serve as isolated microreactors for growth and high-throughput activity-based assays. Non-targeted droplet cultivation of complex microbiomes has enabled recovery of greater microbial diversity than bulk culture, including rare and slow-growing taxa, but lacks the ability to selectively enrich specific functional traits[15–17]. In contrast, only a few studies have reported targeted droplet microfluidic enrichment and characterization from environmental samples[18], and to date, no label-free droplet-based targeted enrichment has been described for proteolytic microorganisms. The strengths of droplet-based microfluidic comprise single-cell encapsulation, precise manipulation, analysis and sorting, using several different detection modalities, amongst the most common: absorbance and fluorescence[19,20]. We previously introduced passive and image-based droplet microfluidic platforms for screening proteolytic microorganisms[21,22], based on the encapsulation of single cells in gelatine droplets and sorting by droplet deformability at higher throughput than earlier deformability-based approaches[23]. These advances highlight the potential of droplet microfluidics to link microbial function to single-cell phenotype and accelerate the discovery of proteolytic species relevant to biogas optimization. In this context, proteolytic consortia remain a key yet insufficiently characterized component of sludge hydrolysis, and this study contributes to ongoing efforts to better understand their composition and role in substrate degradation. Here, we present a droplet microfluidic workflow that integrates two previously developed label-free protocols exploiting changes in the mechanical properties of gelatine droplets to detect and isolate proteolytic microorganisms. This approach enables the enumeration and isolation of proteolytic bacterial cultures originating from single cells encapsulated in picolitre droplets. Following incubation, droplets were first analyzed using the Induced Droplet Ovalization (IDO) method, in which image-based quantification of droplet deformation correlates with microbial proteolytic activity[21]. Subsequently, droplets were sorted at high throughput using a Deformability-based Passive Droplet Sorter (DPDS)[22], enabling the enrichment and downstream taxonomic characterization of proteolytic microcultures. The performance of the proposed droplet-based method was compared with the SMA protocol to evaluate their respective effectiveness in isolating proteolytic consortia from sludge samples. The objectives of this study were to: i. apply high-throughput strategies for functional screening of proteolytic bacteria from complex environmental samples such as sewage sludge; ii. quantify, enrich, and identify proteolytic strains via 16S sequencing; and iii. evaluate droplet-based protocols as scalable, label-free alternatives to conventional approaches such as skim milk agar screening.

## Results and discussion

We cultivated, enriched, and characterized proteolytic strains from sewage sludge, as outlined in Figure 1 (A), and evaluated our novel droplet-based microfluidic workflow against a conventional protocol. Sludge was first resuspended and streaked on solid medium, while in parallel, single cells were encapsulated in gelatine droplets for clonal cultivation. In the traditional SMA assay, community analysis and colony isolation were performed manually, whereas the droplet-based system enabled automated, high-throughput processing. This setup allowed a direct comparison of outcomes between bulk and droplet-based methods.

**Figure 1.**
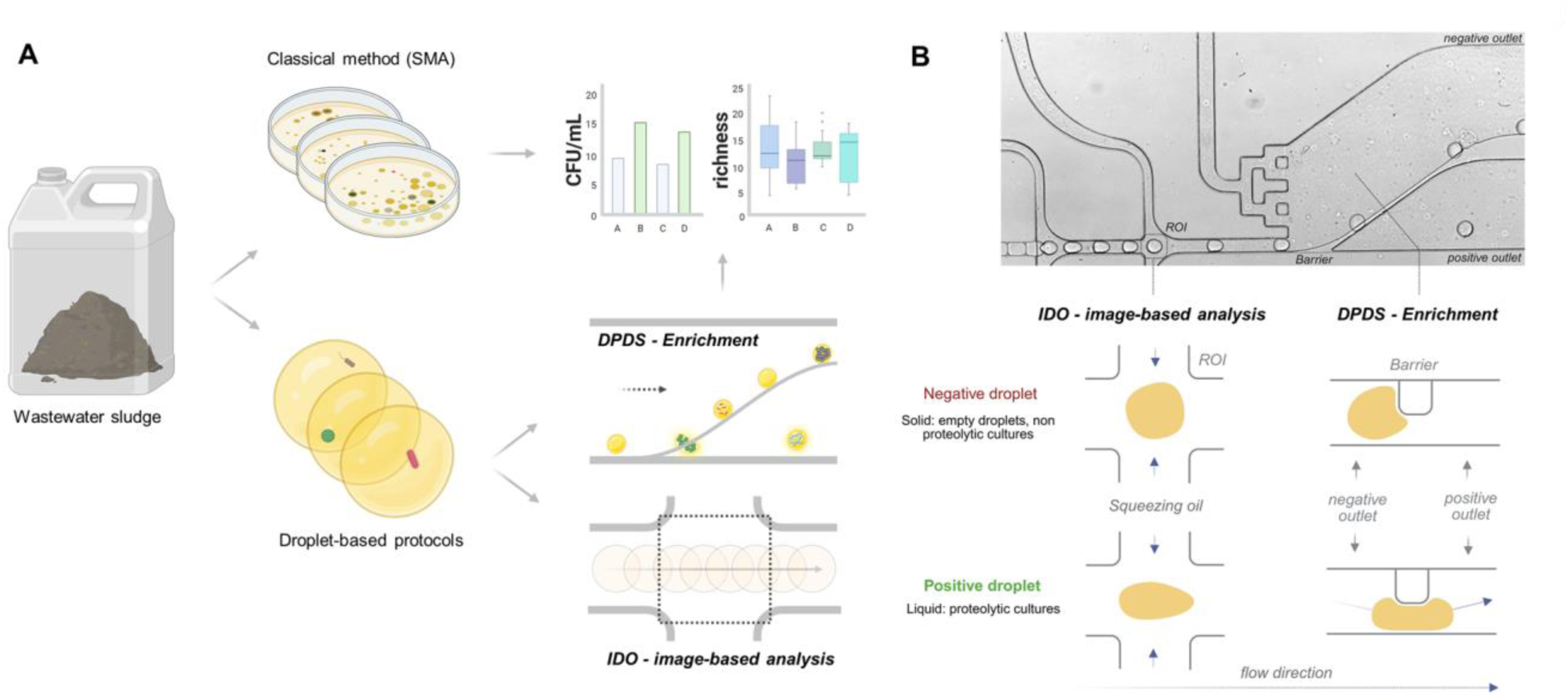
Environmental screening of microbial proteolytic activity. **(A)** Classical screening of sewage sludge was performed by plating microorganisms on skim milk agar (SMA), followed by manual quantification and analysis of colonies displaying proteolytic activity. In parallel, droplet-based microfluidic protocols enabled the encapsulation of single microbial cells into picoliter-sized compartments, where they were cultured and assayed for enzymatic activity in a high-throughput and automated manner. **(B)** In this workflow, gelatine droplets were first characterized using image-based IDO analysis (Induced Droplet Ovalization) to quantify the abundance of proteolytic strains and were then passively sorted via the DPDS device (Deformability-Based Passive Droplet Sorter) for the enrichment of proteolytic strains. This microfluidic protocol offered an effective means to assess microbial activity and diversity in environmental samples, while also serving as a direct benchmark against classical approaches.

We employed two complementary droplet microfluidic protocols to detect proteolytic activity at the single-cell level, Figure 1 (B). In the IDO analysis (Induced Droplet Ovalization), individual cells are encapsulated in gelatine droplets, and microbial protease production alters droplet viscoelastic properties, making the droplets more deformable in a flow-focusing device; these deformations, monitored by automated image analysis, correlate with the proteolytic activity of the encapsulated clonal cultures. Detection of droplet deformability enables label-free, high-throughput identification and enumeration of proteolytic strains[21]. In the DPDS method (Deformability-based Passive Droplet Sorter), gelatine degradation by proteases triggers a solid-to-liquid phase transition, allowing passive microfluidic sorting of liquid droplets (proteolytic strains) from stiffer droplets (empty or non-proteolytic cultures) by flowing droplets over a microbarrier, where only liquid droplets are positively sorted without labels, optoelectronic components, or complex hardware[22]. Finally droplet samples were sequenced to assess taxonomic richness of sewage sludge. Together, these methods enable rapid, high-throughput analysis and enrichment of proteolytic microbial consortia from environmental samples.

### Analysis of the abundance of proteolytic bacteria in sewage sludge using the IDO system

For this study, sewage sludge was selected as the environmental sample for microbial isolation. The sludge sample analyzed in this study was collected in summer (August 2024) and represents a different sample from that used in the previously described sewage sludge analysis conducted for validation of the IDO protocol. The validation study was performed using a winter sample (February 2025) collected from the same biogas facility. In our previous work the sludge sample was already used primarily to validate a droplet-deformability, image-based method for analyzing proteolytic activity [21]. In the present study, we analyzed a different wastewater sample to expand the enrichment protocol [22] by enabling automated and accurate enumeration of positive cultures, followed by their genomic identification, thereby broadening the applicability of a user-friendly droplet-based approach for screening microbial consortia.

As shown in Figure 2, SMA dishes display proteolytic colonies characterised by a transparent halo, whereas colonies lacking the halo were the non-proteolytic strains. Image analysis (IDO) identified positive droplets, which appeared oval-shaped when compressed in the second focusing junction of the sorter device, the DPDS, as shown in Figure 2 (B). In contrast, non-proteolytic and empty droplets remained spherical due to the presence of gelatine within the droplets. The results from the different screenings revealed a substantially higher number of CFU/mL of proteolytic strains detected automatically through IDO analysis compared to the SMA protocol, Figure 2 (C). Interestingly, single-cell encapsulation in droplets also yielded a greater number of non-proteolytic strains, despite the system not being specifically designed for this purpose. These latter strains were enumerated manually; nevertheless, this supports that droplet-based protocols can increase the number of cultivable strains recovered from environmental samples compared with conventional plate-or bulk-based methods, consistent with previous non-targeted cultivation studies reporting improved recovery through droplets[15–17].

**Figure 2.**
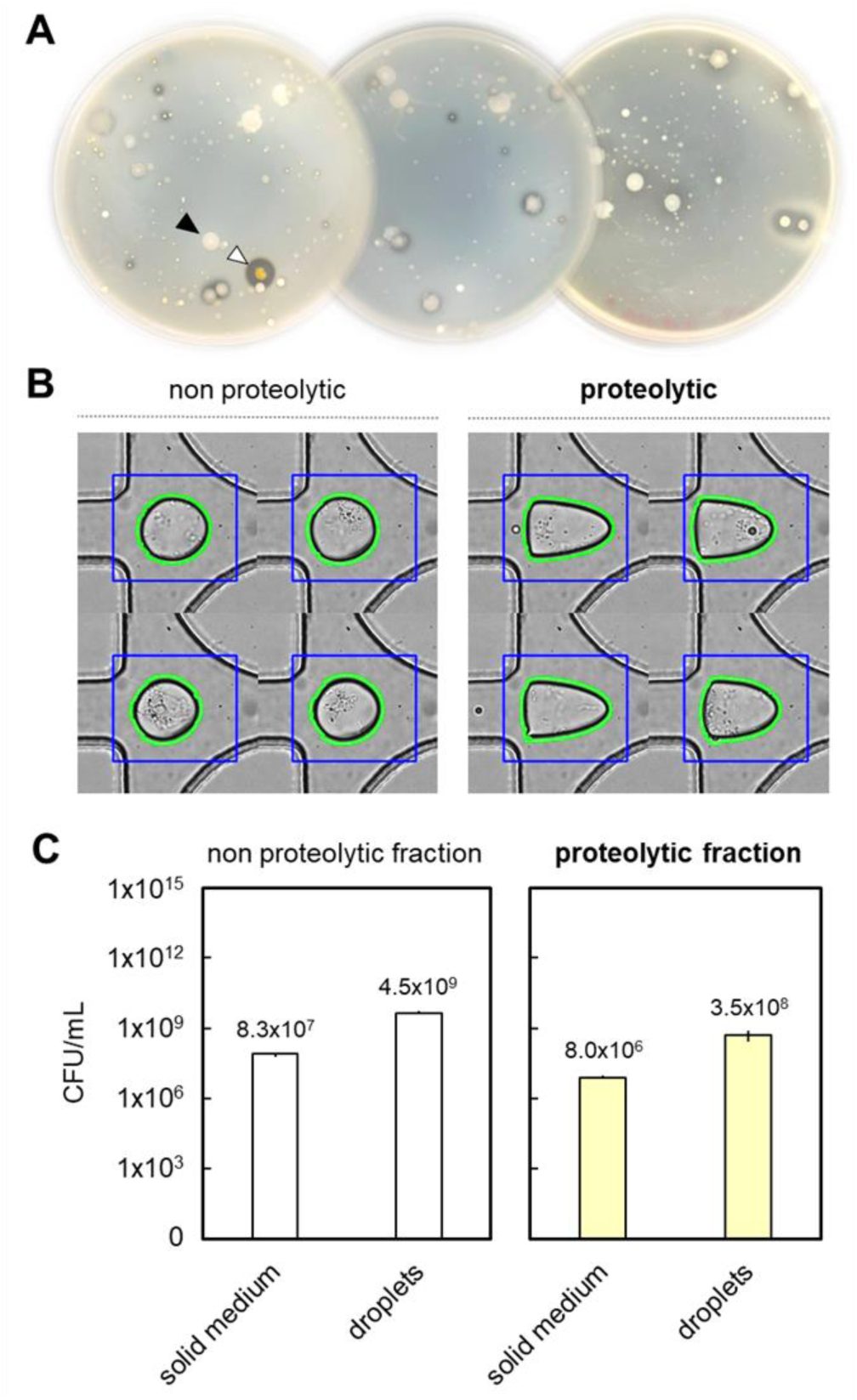
Comparative screening of microbial consortia using solid medium and droplet-based image analysis. (**A**) Representative SMA plates after 96 hours of incubation, showcased phenotypic differentiation of microbial colonies. Colonies surrounded by transparent halos (white arrow) exhibit proteolytic activity visible due to gelatine hydrolysis, whereas colonies without halos (black arrow) correspond to non-proteolytic strains. (**B**) In the image-based analysis (IDO), proteolytic strains exhibit a distinct elongated droplet morphology, resulting from gelatine degradation and the consequent deformation at the DPDS focusing junction. In contrast, droplets carrying non-proteolytic strains retain a spherical morphology. Representative examples of droplets are shown. (**C**) The bar plots illustrate how cultivation in gelatine droplets (λ∼0.2) resulted in approximately two orders of magnitude higher growth for both proteolytic and non-proteolytic strains compared with conventional SMA plating. This highlights the enhanced sensitivity of droplet-based microfluidic analysis relative to traditional solid-medium screening.

### Methodological details of IDO analysis of the sludge sample

Each droplet detected via IDO was calculated as the result of consecutive frames in which a full droplet appeared within a defined region of interest (ROI) in the video[21]. During video analysis, all extracted data were saved in an output file (e.g.,.csv or.xlsx) together with the corresponding screened frames, thereby documenting the detection of droplets within the ROI. In total, 5666 droplets were automatically analysed from video recordings to detect proteolytic colonies, Figure 3 (A, B). IDO analysis was performed using strict area and perimeter thresholds to ensure that only droplets of approximately 100 pL were detected, thereby excluding debris, satellite droplets, and merged droplets from the dataset, as shown in Figure S1. To discriminate between negative solid and spherical droplets (empty or containing non-proteolytic cultures) and positive elongated droplets with proteolytic activity, we applied an aspect ratio threshold of 0.75. Droplets with an aspect ratio below this value were classified as positive. This threshold was determined in our previous study using LB 0.5× droplets with varying gelatine concentrations (0-7.5%), as it corresponds to the solid-to-liquid transition observed between 3% and 4.5% gelatine[21]. The sewage sludge emulsion composition (λ∼ 0.2) is shown in the pie chart. In addition to IDO analysis, proteolytic cultures were enumerated by visual inspection of video recordings to validate the classification. Empty droplets and non-proteolytic cultures, which were not automatically distinguished by IDO analysis, were visually identified and counted. In total, 5666 droplets were classified.

**Figure 3.**
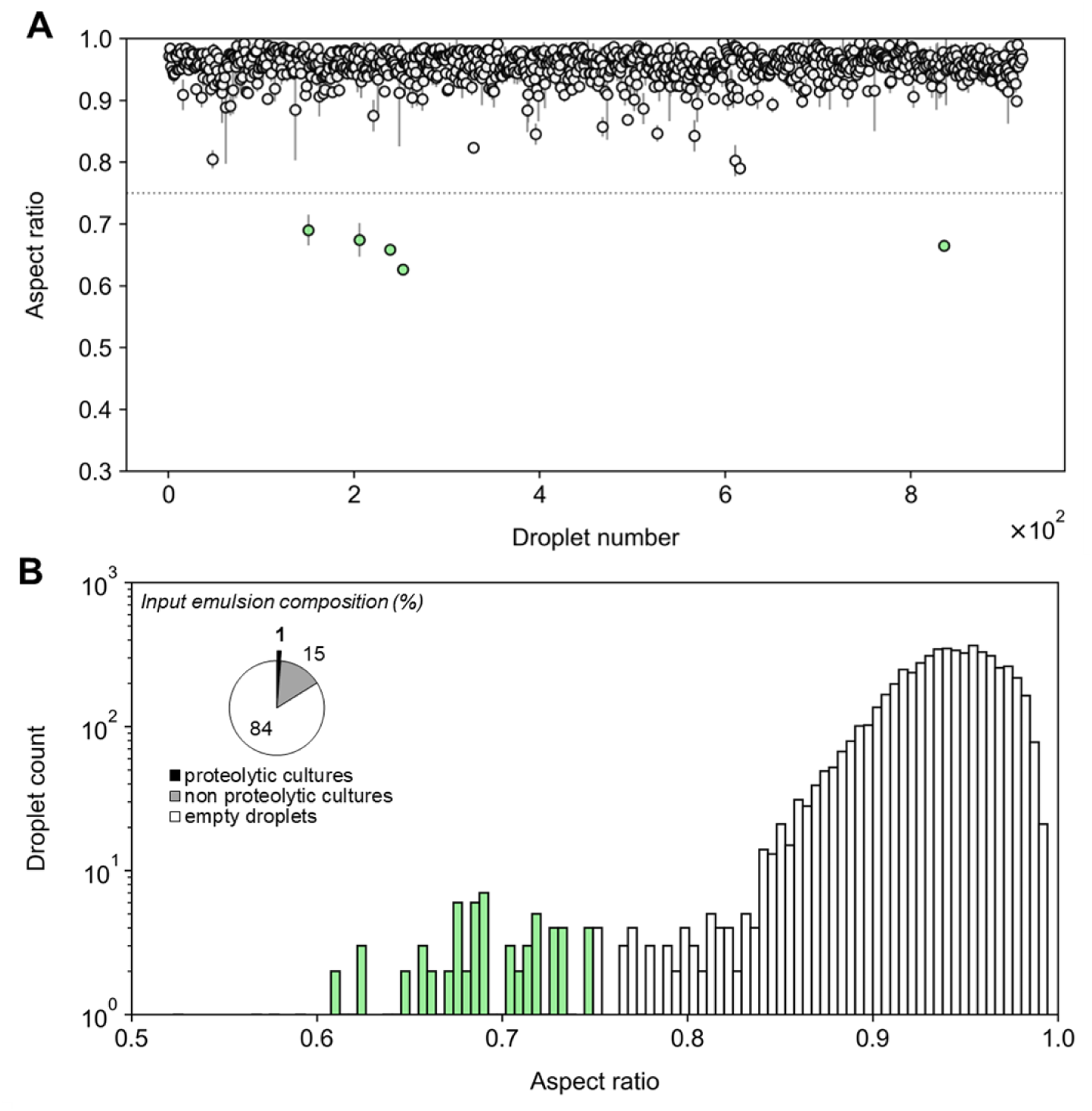
Detection of microbial proteolytic activity. (**A**) The figure summarises the results of the image-based method applied to video recordings from enrichment experiments in the DPDS device. The scatter plot shows the mean aspect ratio of individual droplets. The error bars represent the standard deviation of the aspect ratio calculated across multiple frames for each droplet. Droplets with aspect ratio values below 0.75 (grey dotted line) were classified as liquid, indicating microbial proteolytic activity (green circles). (**B**) The bottom histograms display the aspect ratio distribution of droplets detected from the video recordings, with green bins representing droplets with an aspect ratio below 0.75, which were identified as positive droplets encapsulating individual proteolytic cultures. The emulsion composition (λ ∼0.2) is illustrated in the pie chart, calculated from video recordings and IDO analysis of a total of 5666 droplets.

### Enrichment of proteolytic consortia from sewage sludge

Following the image-based analysis, we aimed to isolate proteolytic bacteria using the DPDS device, which sorts droplets in a passive manner and its performance was previously demonstrated for screening of mock microbial communities[22]. Gelatine droplets with microbial colonies were sorted as shown in Figure 4 (A, B): liquefied droplets containing proteolytic strains were squeezed beneath the barrier and they were directed to the positive outlet, whereas empty droplets and those containing non-proteolytic strains remained solid and could not pass under the barrier. The screening was carried out at a controlled room temperature of 20 °C. Droplet screening was performed for 240 min, which at a throughput of 50 droplets per second (50 Hz) corresponds to approximately 7.2 × 10⁵ sorted droplets. The DPDS outlets were connected via sterile Teflon tubing to sterile Eppendorf tubes in which the droplets were collected. Once the enrichment was completed, the tubes were disconnected from the device, and the emulsions were broken as described in the “Materials and methods” section, resulting in the resuspension of bacteria in physiological solution. The bacterial suspension from merged droplets was divided, with a small fraction (50 µL) analyzed using the SMA test for both positive and negative outlet samples, and the remaining fraction (150 µL) processed for sequencing. SMA analysis of the DPDS outlets indicated a high level of sorter efficiency; however, cultivation on solid media is known to bias the representation of environmental microbial communities[16], this test should be regarded as a qualitative control, Figure 4 (C). A few potential false positives were observed, as non-proteolytic colonies were found in the positive outlet sample. This estimate of approximately 10% false positives in Petri dishes should be interpreted with caution, as growth on solid agar can induce metabolic states that differ from those in liquid droplets. Moreover, the low variability in colony number and morphology observed on SMA among streaked droplet samples suggests that a substantial fraction of strains did not grow, rendering SMA screening poorly informative for accurate quantification at this post-droplet enrichment stage. In contrast, all colonies recovered from the negative outlet lacked proteolytic activity. The limited diversity in morphology and pigmentation of non-proteolytic colonies further indicates reduced variability, likely reflecting the dominance of fast-growing strains on solid media. The output emulsion composition (%), Figure 4 (D), was assessed by streaking droplets sorted from the negative and positive outlets of the DPDS device onto SMA plates.

**Figure 4.**
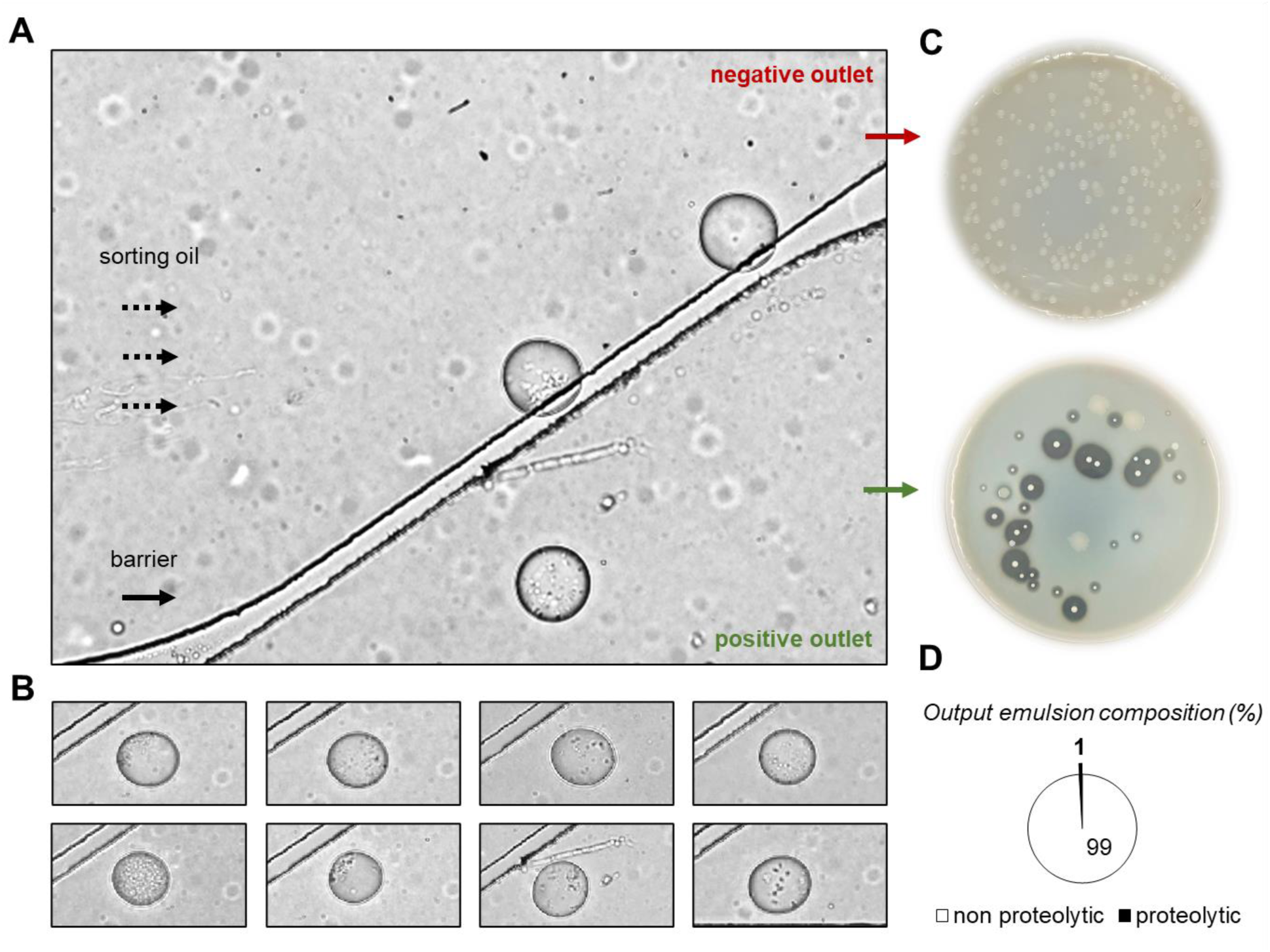
Isolation of proteolytic strains from sewage sludge. (**A**) Individual colonies were sorted simultaneously by the DPDS device: liquefied droplets were squeezed underneath the barrier by the laminar flow of oil and directed to the positive outlet. Empty droplets and those containing non-proteolytic strains, which remained solid and could not pass under the barrier, were directed to the negative outlet. (**B**) Snapshots of positive droplets during sorting were captured during the enrichment. (C) Droplets collected from both outlets were further characterised using SMA to assess sorter efficiency. Colonies recovered from positive droplets were predominantly proteolytic, except for a few possible non-proteolytic strains. In contrast, all colonies from negative droplets were non-proteolytic. (D) The output emulsion composition (%) was calculated based on the CFUs screened using the SMA protocol following DPDS enrichment.

### Droplet-based cultivation of sewage sludge-derived microorganisms enhances microbial diversity and abundance

#### Validation of sorting of proteolytic cultures from a binary mock community

To assess the performance of the microfluidic enrichment performed with DPDS device, we sequenced the V4-V5 region of the 16S rRNA gene on an Illumina NovaSeq platform from a binary mock community composed of *Pseudomonas aeruginosa* (λ∼0.01), a highly proteolytic reference strain previously isolated from the sewage sludge, and *Escherichia coli* (λ∼0.1) as the non-proteolytic counterpart, with λ represents the mean cell occupancy per droplet under Poisson loading conditions. Based on the emulsion composition and the screening duration, the estimated total number of droplets collected from the DPDS outlets was approximately 3.6 × 10⁵ droplets (100 pL each) during a 2-hour screening (7.2 × 10³ s at 50 Hz). The sequencing analysis revealed that the DPDS device effectively enriched the proteolytic *Pseudomonas aeruginosa*. In fact, approximately 95% of reads assigned to *Pseudomonas* and *Escherichia-Shigella* were detected at minimal relative abundance (5%), indicating a very low rate of false positives. Conversely, the negative outlet showed 4 ASVs distributed across three genera, with a predominance of *Escherichia-Shigella* (75%) and approximately a 25% relative abundance of false negatives represented by *Pseudomonas,* Figure 5 (A). The relatively high rate of false negatives may be attributed to phenotypic variation. In previous experiments, a gradual loss of proteolytic activity in *P. aeruginosa* was observed during cultivation in gelatine droplets, suggesting instability of the expressed phenotype over time. Similar effects are common in conventional screening approaches, where it is recommended to revive cultures from glycerol stocks after several passages to minimise phenotypic drift. This effect may be further amplified in droplet-based assays, given the high-throughput nature of the process and the large number of screened events. The *P. aeruginosa* strain used in this study was isolated from sludge by selecting a single positive colony on SMA[24–26]. As an environmental isolate, it is likely subject to greater phenotypic variability than a stabilised *E. coli* line, leading to subpopulations that lose proteolytic activity. Consequently, some *P.aeruginosa* droplets remain solid, preventing their correct sorting under the optimized conditions. By contrast, the laboratory strain we tested *E. coli* DH5α exhibits markedly higher genetic and phenotypic stability. Such stability renders it considerably less prone to phenotypic variation than wild-type or environmental isolates, which often display heterogeneous behaviour under stress or during prolonged cultivation. Overall, these results confirm that sorting based on the positive outlet reliably enriches proteolytic strains, demonstrating the effectiveness of the DPDS device in selecting the desired phenotypes despite some unavoidable phenotypic variability.

**Figure 5.**
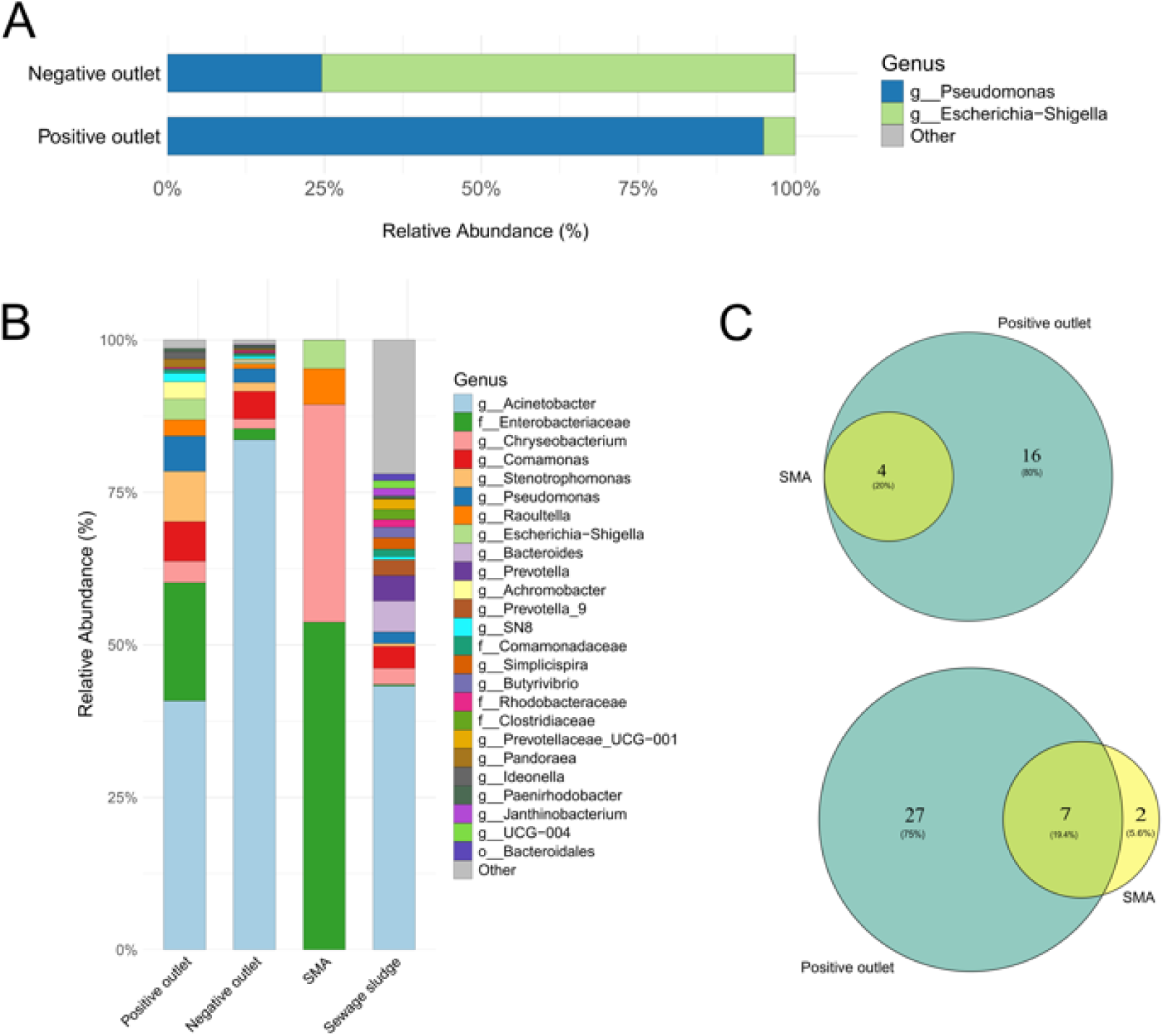
Comparison of microbial diversity recovered by microfluidic droplet-based enrichment and SMA plate culturing. (**A**) Genus-level taxonomic composition of the most abundant genera in the positive and negative droplet sorter outlets for mock consortia experiment. All remaining taxa are grouped as “Other”. (**B**) Genus-level taxonomic composition of the most abundant genera in original sludge, SMA plate isolates, and the positive and negative droplet sorter outlets. All remaining taxa are grouped as “Other”. (**C**) Venn diagrams showing the overlap of genera (up) and ASVs (down) recovered by SMA plating and the positive microfluidic outlet.

#### Droplet-based screening of sewage sludge compared with traditional protocol

After validating the reliability of the device output, we assessed its ability to recover proteolytic strains from a complex sewage sludge sample by comparing its performance with conventional SMA cultivation method. The V4–V5 region of the 16S rRNA gene was sequenced (Illumina NovaSeq) from: (i) droplets collected from both DPDS sorter outlets, (ii) proteolytic colonies isolated on SMA medium, and (iii) the original sewage sludge sample. The sludge was resuspended as described in the “Materials and methods” section; one aliquot was sequenced directly, while the remaining suspension was used for droplet encapsulation and SMA-based screening. The estimated number of droplets collected through the positive outlet of the DPDS was calculated based on the emulsion composition and the screening duration. During a 4-hour screening (1.44 × 10⁴ s at 50 Hz), approximately 7.2 × 10⁵ droplets (100 pL each) were generated. Of these, 1.3% were positive (∼9,360 droplets), corresponding to a final collected volume of approximately 0.9 μL. In contrast, conventional screening on SMA medium yielded only a limited number of proteolytic colonies. After incubation, a total of 32 colonies exhibiting proteolytic activity, as indicated by transparent halos, were hand-picked from multiple SMA plates and sequenced.

To assess whether the microfluidic device recovered a higher diversity of proteolytic species than SMA plates, we compared the two samples using rarefaction–interpolation analysis[27] based on Hill numbers (q = 0, 1, 2) with 95% confidence intervals[28]. Diversity extrapolation curves showed that the positive outlet exhibited substantially higher ASV richness than SMA cultivation, Figure S2 (A). This increase was driven by a broader taxonomic composition, with droplet-derived communities enriched in rare and low-abundance taxa that were largely absent from plate-derived samples. Rank–abundance distributions confirmed higher evenness in droplet samples, whereas SMA plates were dominated by a few fast-growing taxa, Figure S2 (B). This pattern is consistent with the known advantages of droplet microfluidics, where the physical compartmentalization of individual cells minimizes competitive interactions and promotes the growth of rare and slow-growing microorganisms[16,29,30].

A closer examination of the taxonomic composition reveals striking differences between the two cultivation approaches. Sequencing of the sludge sample revealed 43.28% of reads assigned to *Acinetobacter*, followed by *Bacteroides* (5%); members of the *Prevotella* genus such as *Prevotella* (4.1%) and *Prevotella-9*; *Comamonas* (3.6%); *Chryseobacterium* (2.6%); and several other genera, each representing <2% relative abundance as reported in Figure 5 (B) and Table S1. This pattern is consistent with previous studies reporting *Acinetobacter* as a frequent and often dominant genus in wastewater treatment plants, particularly in sewage sludge[31–33]. Its relative abundance, however, is known to vary depending on wastewater source, plant type, operational conditions, and seasonal fluctuations, resulting in distinct microbial profiles across sludge samples[34–36]. While the original sewage sludge contained high diversity, SMA plates recovered only four genera: Enterobacteriaceae family (53.75%), *Chryseobacterium* (35.66%), *Raoultella* (5.09%), and *Escherichia–Shigella* (4.7%). Notably, the dominant genus of the sludge, *Acinetobacter,* was completely absent from the SMA plate, Figure 5 (B). All detected taxa belong to bacterial groups with documented proteolytic capabilities, including the production of extracellular and/or cell-associated proteases involved in the degradation of proteinaceous substrates[37–39]. Droplet-based cultivation and sorting, in contrast, preserved sludge diversity more faithfully: both sorter outlets were dominated by Acinetobacter (40.8% positive, 83.6% negative), recovering 20–23 genera. In the positive outlet, following *Acinetobacter*, reads were assigned to the Enterobacteriaceae family (19.38%), *Stenotrophomonas* (8.2%), *Comamonas* (6.4%), *Pseudomonas* (5.8%), and *Chryseobacterium* (3.5%). Although a small fraction of genera may represent false positives, the majority are known proteolytic taxa (e.g., *Chryseobacterium*, *Pseudomonas*, *Acinetobacter*, *Staphylococcus*)[40–43]. The occasional detection of non-proteolytic taxa, such as *Microvirgula* (0.15%), and members of Rhodobacteraceae (0.25%), aligns with observations from the mock community, where false positives accounted for only ∼5% of reads, Figure 5 (A). Crucially, the comparative analysis of taxa recovered by SMA plate selection and microfluidic enrichment revealed that the four genera isolated on the SMA plate were entirely recapitulated in the positive microfluidic output. The observed high level of agreement was maintained at higher taxonomic resolution, with 7 out of the 9 ASVs detected on the plate being successfully captured by the microfluidic device, as shown in Figure 5 (C).

Droplet-based enrichment (positive outlet only) enhanced low-abundance genera with relevant functional potential compared to conventional cultivation. In the sludge sample, most detected taxa (87.9%) were below <1% relative abundance (Table S2). Ten low-abundance genera were recovered in droplets and/or plate cultures, six of which exceeded 1% abundance after enrichment (Table S3). Notably, all ten were detected in the positive droplet outlet, whereas SMA screening recovered only 2 (*Raoultella* and members of the *Enterobacteriaceae* family), highlighting the advantage of droplet cultivation for accessing rare taxa. Several enriched genera include strains with extracellular proteolytic activity or high metabolic versatility. *Stenotrophomonas maltophilia* produces multiple secreted proteases, including the extracellular serine proteases StmPR1/2/3, which are involved in substrate hydrolysis and virulence and show strong proteolytic activity in culture assays [44]. Members of the *Enterobacteriaceae* family, such as *Raoultella*, encode diverse protease systems (e.g. Clp proteases and other peptidases) supporting protein turnover and nutrient acquisition [45]. *Pandoraea* species, mainly environmental bacteria and opportunistic pathogens, display metabolic traits linked to biodegradation and environmental persistence, although their extracellular proteases remain poorly characterised and merit further study [46]. The enrichment of these taxa in droplets, particularly above 1% relative abundance, highlights the utility of droplet micro-compartmentalization for accessing taxa with *in situ* metabolic capacity that might otherwise be outcompeted in culture.

The negative outlet also yielded a substantial number of genera, with *Comamonas* as the second most abundant (4.5%), followed by *Pseudomonas* (2.2%), Enterobacteriaceae (1.8%), *Chryseobacterium* (1.6%), and *Stenotrophomonas* (1.6%); all other genera were present at <1% relative abundance, Figure 5 (C) and Table S1.

Overall, these results demonstrate that droplet-based microfluidic cultivation enables a more comprehensive recovery of microbial diversity than conventional plate methods, substantially increasing the capture of proteolytic species that are otherwise underrepresented. By enriching for low-abundance, potentially slow-growing or hard-to-cultivate taxa, some of which are known to exhibit extracellular proteolytic or other hydrolytic activities, droplet cultivation broadens both the phylogenetic and functional scope of cultivable isolates. While conventional SMA screening often fails to capture the full breadth of community complexity, microfluidic droplet technology significantly expands the discovery landscape, enabling the identification of rare microbial strains with specialized enzymatic profiles for protein degradation and other extracellular activities.

## Conclusion

Droplet microfluidics offers several advantages for conducting biochemical assays within stable microdroplets, including reductions in volume and reagent consumption, enabling screening with an unprecedented level of throughput, with current protocols able to screen up to 10⁸ samples per day[47].

Moreover, droplet-based systems facilitate single-cell analyses by allowing the cultivation and screening of microcultures originating from individual cells, thereby enabling the detection of rare and slow-growing taxa that are often missed by conventional approaches[16]. Droplet-based cultivation and sorting enable high-throughput screening of environmental samples[18], yet proteolytic strains remain largely unexplored despite their relevance, for example, in substrate hydrolysis. Our workflow provides a robust framework for the functional screening and taxonomic characterization of proteolytic microbes from sewage sludge, supporting a comprehensive understanding of their composition. In this study, we demonstrate that droplet-based microfluidic screening provides a powerful and scalable strategy for the functional analysis (IDO) and enrichment (DPDS) of proteolytic microbial consortia from environmental samples such as sewage sludge. By combining single-cell encapsulation in gelatine microcompartments, label-free image-based detection of proteolytic activity, and passive droplet sorting of proteolytic cultures, we establish a comprehensive workflow that overcomes several inherent limitations of conventional bulk methods. This approach achieves substantially higher throughput and, more importantly, enables the screening of a larger number and greater diversity of microbial strains.

Importantly, when benchmarked against other droplet-based HTS workflows, our approach offers substantially greater practical simplicity. Existing droplet microfluidic platforms for environmental HTS typically rely on fluorescence-activated droplet sorting (FADS) to isolate proteolytic strains[48]. Although these methods achieve higher throughput than our protocols, they require specialised substrates or fluorogenic reporters as well as complex optical systems[49]. In comparison, the label-free, image-based detection coupled with passive sorting eliminates reporter’s use and lowers system complexity while retaining lower but still high throughput figures.

We therefore proposed a direct comparison of our workflow with skim milk agar screening, a widely used in-bulk method for the detection of proteolytic activity. The results show that solid-medium cultivation substantially underestimates both the abundance and diversity of proteolytic microorganisms. In contrast, droplet-based systems recovered a richer community, including genera that were absent or severely underrepresented on plates. Droplet enrichment captured all proteolytic genera identified by classical cultivation and additionally enabled the isolation of a substantially broader range of known or putatively proteolytic taxa. The 16s rRNA sequencing of sorted droplet populations further demonstrated that microfluidic enrichment yields communities that more closely reflect the original sludge microbiome, while selectively enriching for proteolytic phenotypes. By incorporating high-throughput sequencing directly after droplet sorting, we gained deeper insights into the ecological and biotechnological potential of these communities than approaches where sequencing follows traditional cultivation or bulk extraction, in which only a fraction of what grows in liquid droplets can be recovered on solid media[48]. Future developments will focus on integrating the present high-throughput droplet screening platform with droplet deposition systems to enable direct transfer of sorted microcultures into individual wells[50]. Such coupling would bridge single-cell, in-droplet cultivation and selection with automated downstream upscaling[51].

Overall, this work further expands droplet microfluidic methods, establishing them as a robust platform for functional HTS of microorganisms. The ability to selectively enrich active proteolytic microorganisms opens new opportunities for analysing hydrolytic microbial communities and recovering strains with potential applications in biomass hydrolysis, bioremediation, and industrial enzyme discovery. Our approach provides a general framework for linking microbial function to community composition in complex ecosystems. Furthermore, our methods can be readily adapted to diverse screening campaigns, provided that the solid-to-liquid transition of droplets can be exploited to trigger the deformability-based sorting, for example to detect agarolytic activity.

## Materials and Methods

### Resuspension of the environmental sample

Sewage sludge was sampled in August 2024 from a biogas facility in Wołomin, Poland, and stored at 4°C in a 5 L plastic container. Approximately 25 mL were transferred into 50 mL centrifuge tubes and centrifuged at 2000 rpm for 2 min to sediment particulate matter and concentrate the sample. One milliliter of sediment was resuspended in 100 mL sterile physiological saline in a 500 mL Erlenmeyer flask and incubated at 25°C with shaking at 250 rpm for 12 hours to facilitate sample resuspension. The suspension was subsequently passed through a 40 µm cell strainer mesh filter (VWR) to remove coarse particles, and the filtrate was collected for downstream experiments.

### Gelatine droplet generation and cultivation of microbes

Preliminary screening of sewage sludge on LB agar was used to estimate the CFU/mL of the sample. After overnight shaking in physiological solution, an appropriate volume of the cell suspension was transferred to a 1.5 mL microcentrifuge tube to achieve the desired cell-to-droplet ratio (λ) in the resulting emulsion. Cells were collected by centrifugation (5000 × g for 5 min), the supernatant was discarded, and the cell pellet was thoroughly resuspended in gelatine medium. The medium was prepared in ddH₂O and consisted of 5 g/L NaCl, 2.5 g/L yeast extract, 5 g/L tryptone, and 75 g/L gelatine. The solution was then sterilized by autoclaving. Droplet emulsions were generated using a flow-focusing droplet generator with an oil phase containing 5% RAN 008-Fluorosurfactant at 25°C (RAN Biotechnologies). The emulsions were collected in a droplet chamber and incubated at 40°C for 96 hours. A peristaltic pump supplied oxygen dissolved in the oil to the bacteria inside the droplets, while preventing air bubbles from entering the chamber[21,22]. Detailed protocols for droplet generation and microbial cultivation are available in the supplemental Information (SI), Figure S3.

### Characterization of proteolytic consortia in the input samples

Sludge was streaked on Petri dishes on the same day as droplet generation by inoculating 100 µL of different serial dilutions (10⁻², 10⁻³, 10⁻⁴) of the suspension, with 6 replicates per dilution, on skim milk agar (SMA). The SMA medium was prepared and autoclaved with the following composition: yeast extract 1 g/L, tryptone 4 g/L, agar 15 g/L, and skimmed milk powder 30 g/L. Plates were incubated at 40 °C and positive colonies were manually picked. The sample was resuspended in a 1.5 mL Eppendorf tube containing 200 µL of physiological solution and stored at-20 °C until sequencing.

### Image-based analysis of droplets encapsulating proteolytic cultures

After incubation, the emulsion was reinjected from the incubation chamber into the DPDS device for image-based analysis. IDO analysis was performed using an optimized Python script that processes high-speed video to extract shape-based features of droplets flowing through the flow focusing section of the device. The script applies background subtraction and contour detection, followed by morphometric analysis to quantify descriptors such as droplet area, perimeter, and aspect ratio. Annotated video frames and quantitative measurements are saved, allowing downstream analysis of droplet deformation as a proxy for microbial proteolytic activity.

### Enrichment of droplets and characterization of proteolytic consortia

Passive sorting of proteolytic microbes was performed in the DPDS device. At the end of sorting, the tubing connected to the outlets were flushed with sterile, filtered HFE-7500 oil. The emulsions were broken by adding 20% (v/v) 1H,1H,2H,2H-perfluoro-1-octanol (PFO, Alfa Aesar) and 200 μL of sterile 0.9% NaCl solution. The biphasic mixtures were vortexed for 90 s and centrifuged for 60 s at 2000 rpm to promote phase separation. The aqueous phases from the outlets were collected, stored for sequencing, and streaked in triplicate on SMA. Plates were incubated at 40°C and examined visually.

### DNA Extraction, DNA Amplification, and Sequencing

DNA was extracted from six samples. These included (i) a sewage sludge sample, resuspended as described in the previous section; (ii) four droplet-derived samples; and (iii) one pooled sample consisting of 32 proteolytic colonies isolated using the skim milk agar (SMA) method. The four droplet samples were obtained after enrichment and sorting with the DPDS device, applied either to the environmental sludge or to a mock community composed of *Pseudomonas aeruginosa* (previously isolated from the same sludge) and *Escherichia coli* as a non-proteolytic control For each sample, DNA was extracted from 100 µL of starting material using the DNeasy Blood & Tissue Kit (Qiagen), following the manufacturer’s protocol for gram-negative bacterial cultures with an initial modification. Due to the low amount of cells in the droplet-derived samples, no pelleting step was performed to avoid material loss; instead, reagents were added directly to the liquid culture aliquots.

Extracted DNA was amplified using the universal prokaryotic primers 515F-Y (5′-GTGYCAGCMGCCGCGGTAA-3′) and 926R (5′-CCGYCAATTYMTTTRAGTTT-3′), targeting the V4–V5 hypervariable region of the 16S rRNA gene[52]. PCRs were performed in triplicate using Phusion High-Fidelity DNA Polymerase (Thermo Fisher). Each 25 µL reaction contained 1 ng of template DNA, 1× Phusion HF buffer, 0.25 μM barcoded forward and reverse primers, 0.02 U/µL polymerase, and 200 μM of each dNTP. Cycling conditions consisted of an initial denaturation at 98 °C for 4 min, followed by 20 cycles of 98 °C for 20 s, 50 °C for 30 s, and 72 °C for 10 s, with a final extension at 72°C for 5 min. Triplicate reactions were pooled to minimize intra-sample variability and to obtain sufficient amplicon quantity, then purified using AMPure XP magnetic beads (Beckman Coulter).

Purified DNA was quantified using a NanoPhotometer NP80 (Implen), and fragment size distribution was assessed via 1% agarose gel electrophoresis. Samples were then sent to the University of Warsaw Sequencing Facility (CeNT Genomics Core Facility) for library preparation and sequencing on the NovaSeq 6000 next-generation sequencing platform (Illumina).

### Sequencing and analysis

Sequence quality was assessed using FastQC[53], and reads were processed in QIIME2. Primer sequences were removed with Cutadapt[54], and denoising was performed using the DADA2 pipeline to infer amplicon sequence variants (ASVs), including paired-end merging and chimera removal (Table S4)[55]. Taxonomic assignment of the V4–V5 16S rRNA gene region was carried out using a Naive Bayes classifier trained on the SILVA 138.1 database and tailored to the 515F-Y/926R amplicon region[56,57]. Downstream analyses were conducted in R (version 4.2.0) using the *phyloseq* package[58] in RStudio 2023.09.0[59]. Rarefaction analyses indicated sufficient sequencing depth for all samples, Figure S4. Prior to downstream analyses, ASVs classified as Eukaryota, chloroplasts, or mitochondria were removed. Rarefaction–extrapolation and rank–abundance analyses were performed on this unfiltered ASV table using the *iNEXT* package[27] to preserve the contribution of low-abundance taxa and accurately assess community structure and sampling completeness, Figure S2. In contrast, taxonomic composition analyses were conducted after filtering low-abundance ASVs to reduce the influence of spurious variants and improve the robustness of taxonomic profiles. Relative abundances were used for comparative taxonomic analyses.

When exact taxonomic annotation was not achievable, sequences were assigned to the lowest reliably supported taxonomic rank for bacteria. Due to the intrinsic limitations of Illumina short-read sequencing, which provides insufficient phylogenetic resolution for deep-level discrimination, confident taxonomic assignment below the genus level was not feasible. Accordingly, strain-level classifications were avoided, as such assignments would be unreliable or inconsistent across samples.

## Supplemental Information

The supplemental Information includes an overview of the bioinformatic workflow and sequencing quality metrics; the distribution and relative abundances of ASVs; analyses of low-abundance (<1%) genera across samples; and the occurrence and enrichment of rare genera from sludge samples under different experimental conditions (Tables S1–S4). Additional figures and videos are also provided.

## Data availability

The research data used to obtain the results presented in the publication have been deposited in the University of Warsaw repository at the following address doi:10.58132/YZ3CEF. NGS data have been deposited in the EMBL-EBI European Nucleotide Archive (ENA) under the Project ID PRJEB107907

## Acknowledgements

This research was funded by the TEAM-NET programme of the Foundation for Polish Science no. POIR.04.04.00-00-14E6/18-00 as a part of Measure 4.4 of the 2014–2020 Smart Growth Operational Programme, EU, and by National Science Centre, Poland (grant SONATA BIS no. 2023/50/E/ST4/00545). Research infrastructure used in the project was co-funded by the “*Excellence Initiative – Research University (2020-2026)*” programme via Action I.4.2 “*Fund for the Renovation and Development of Research Infrastructure*”. During the preparation of this work, the authors used ChatGPT in order to improve language and readability. All content was subsequently reviewed and edited by the authors, who take full responsibility for the content of the published article. The schemes of the workflow shown in Figure 1 were prepared with BioRender.com. We would like to thank Anna Karnkowska and her team from the Institute of Evolutionary Biology for their support and for providing access to tools, methodologies, and expertise that were instrumental to this study.

